# Quercetin Enhances Brown Adipose Tissue Thermogenesis and Improves Glucose-Lipid Metabolism via the COX2-PGE2-EP4-UCP1 Signaling Axis

**DOI:** 10.64898/2026.05.15.725368

**Authors:** Zengqiang Xu, Xiujuan Gao, Jian Sun, Mengxin Jiang, Jiahao Zhu, Yang Geng, Shangzhen Jin, Yang Wang, Yingjiang Xu

**Affiliations:** Binzhou Medical University Hospital, 661 Huanghe 2nd Rd, Binzhou City, 256603, P.R.China; Shandong Medical and Pharmaceutical University, 522 Huanghe 3rd Rd, Binzhou City, 256603, P.R.China

**Keywords:** Brown adipogenesis tissue, Thermogenesis, Quercetin, Cox2

## Abstract

The activation of thermogenesis in brown adipose tissue (BAT) represents a pivotal target for ameliorating disorders of glucose and lipid metabolism. This study sought to elucidate the regulatory effects of quercetin on thermogenesis and glucose-lipid metabolism within brown adipocytes, alongside its underlying molecular mechanisms. The findings demonstrated that quercetin markedly upregulated the expression of uncoupling protein 1 (UCP1), a critical thermogenic protein in brown adipocytes, thereby enhancing cellular thermogenic capacity and effectively mitigating glucose and lipid metabolism disorders. Subsequent mechanistic investigations confirmed that quercetin activated the COX2-PGE2-EP4-UCP1 signaling axis by augmenting the stability of cyclooxygenase 2 (COX2) protein, thus mediating its thermogenic-promoting and metabolism-improving effects. This study identifies quercetin as a potential therapeutic agent for the improvement of glucose and lipid metabolism disorders, uncovers a novel molecular mechanism through which quercetin regulates brown adipocyte thermogenesis, and provides a theoretical and experimental foundation for the application of quercetin in the prevention and treatment of obesity and related metabolic diseases.

## Background

Obesity and its associated metabolic disorders, such as type 2 diabetes mellitus, dyslipidemia, and non-alcoholic fatty liver disease, have emerged as significant global public health challenges, characterized by increasing prevalence and posing substantial threats to human health^1,2^. Additionally, these conditions impose considerable economic burdens on society. In contrast to white adipose tissue (WAT), which primarily functions in energy storage, brown adipose tissue (BAT) is a specialized thermogenic tissue capable of dissipating energy as heat through non-shivering thermogenesis (NST). This process is crucial for maintaining energy balance and glucose-lipid homeostasis within the body^3^. The core effector molecule of BAT thermogenesis is uncoupling protein 1 (UCP1), which is uniquely expressed in the inner mitochondrial membrane of brown adipocytes. UCP1 facilitates the uncoupling of oxidative phosphorylation from ATP synthesis, thereby converting chemical energy into heat. The expression level and activity of UCP1 are directly correlated with the thermogenic capacity of brown adipocytes^4^. Consequently, the activation of BAT thermogenesis and the enhancement of UCP1 expression are considered promising strategies for the prevention and treatment of obesity and its related metabolic disorders.

Quercetin (QUE), a natural flavonoid widely existing in fruits (such as apples, grapes), vegetables (such as onions, broccoli) and traditional Chinese medicines (such as Sophora japonica, Ginkgo biloba), has attracted extensive attention due to its diverse biological activities and high safety^5^. Previous studies have confirmed that quercetin exerts significant regulatory effects on glucose and lipid metabolism, including reducing blood glucose levels, improving insulin resistance, lowering serum triglyceride and cholesterol levels, and inhibiting adipocyte hypertrophy^6–8^. In addition, quercetin also has anti-oxidative, anti-inflammatory and anti-apoptotic properties, which can alleviate metabolic inflammation induced by obesity^9–13^. However, despite the clear regulatory effect of quercetin on glucose and lipid metabolism, its specific regulatory effect on brown adipocyte thermogenesis (especially the regulatory effect on UCP1 expression and activity) and its underlying molecular mechanism remain unclear, which limits the in-depth development and clinical application of quercetin in the field of metabolic disorders.

Building on the previously discussed research background and the identified gaps in the literature, the primary research objective of this study is methodically structured to follow a logical sequence: initially verifying the effect, followed by target screening. The study first seeks to determine the thermogenic activation effect of quercetin on brown adipocytes, specifically examining whether quercetin can enhance UCP1 expression, increase the thermogenic capacity of brown adipocytes, and improve disorders related to glucose and lipid metabolism. Subsequently, omics technologies are utilized to identify potential molecular targets involved in quercetin’s regulation of thermogenesis. Upon identifying COX2 as a critical target through omics screening, the research further investigates the molecular mechanisms, with a focus on COX2 protein stability and the PGE2-EP4-COX2-UCP1 signaling axis.This investigation aims to ascertain whether the thermogenic and metabolic advantages of quercetin are facilitated by the stabilization of the COX2 protein and the activation of the related signaling pathway. The study endeavors to elucidate the emerging regulatory role of quercetin in maintaining metabolic homeostasis, to uncover novel molecular mechanisms by which quercetin influences brown adipocyte thermogenesis through omics-based target screening, and to provide both a theoretical framework and empirical evidence for the potential development of quercetin as a natural therapeutic agent for the prevention and treatment of obesity and related metabolic disorders.

## Results

### Result 1: QUE Alleviates Obesity and Metabolic Disorders Induced by a High-Fat Diet

To examine the impact of QUE on obesity and associated metabolic disorders, 6-week-old male C57BL/6J mice were subjected to a HFD and received intraperitoneal injections of QUE at doses of 10 mg/kg, 50 mg/kg, or 100 mg/kg, or saline, over a period of 9 weeks (Figure 1A). Initially, the effects of varying doses of QUE on body weight were assessed. The body weight trajectory, encompassing all three doses (10, 50, and 100 mg/kg, Figure 1B), demonstrated that all QUE-treated groups experienced a significant reduction in body weight gain compared to the HFD group. Notably, the 50 mg/kg dose of QUE exhibited the most substantial inhibitory effect on body weight gain. Consequently, the 50 mg/kg dose was selected as the optimal dose for subsequent experiments. In the body weight trajectory using only the 50 mg/kg dose (Figure 1C), QUE treatment continued to significantly suppress body weight gain, leading to a notably reduced final body weight (Figure 1C) and a significantly lower area under the body weight curve (AUC, Figure 1C). Consistently, QUE intervention at 50 mg/kg significantly reduced the tissue weights of BAT,iWAT,eWAT and liver (Figure 1D).

**Figure 1.**
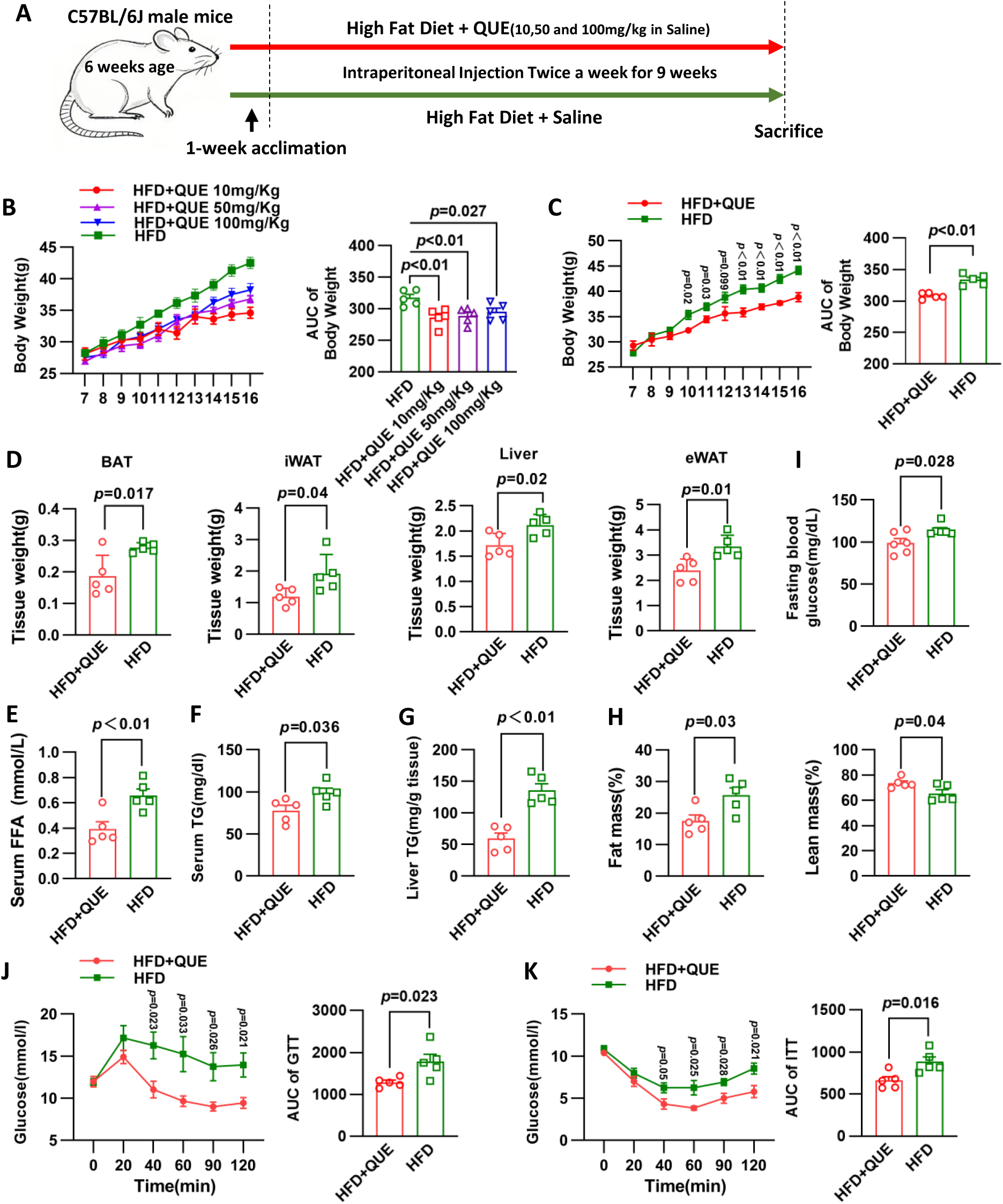
QUE ameliorates high-fat diet-induced obesity and systemic glucose-lipid metabolic disorders. (A) C57BL/6J male mice fed on a high-fat diet. HFD was fed to mice at 6 weeks of age and intraperitoneally injected with QUE (10 mg/kg, 50 mg/kg, or 100 mg/kg) or saline for 9 weeks. (B) Body weight of mice in (A) during high-fat diet feeding and AUC of the left. N = 5 mice in each group. (C) Body weight of mice Treated QUE(50mg/kg) during high-fat diet feeding and AUC of the left. n = 5 mice in each group. (D) Tissue weight of BAT, iWAT, eWAT and Liver in high-fat diet treated mice. N = 5 mice in each group. (E-G) Serum FFA(E), Serum triglyceride(F), Liver triglyceride(G) concentrations were assayed in high-fat diet treated mice. n = 5 mice in each group. (H)Assessment of lean and fat mass by quantitative nuclear magnetic resonance of high-fat diet treated mice. n = 5 mice in each group. (I)Fasting blood glucose of high-fat diet treated mice. n = 5 mice in each group. (J)Left, GTT in mice after 9 weeks of HFD feeding. Right, AUC was used to quantify the GTT results. n = 5 mice in each group. (K)Left, ITT in mice after 9 weeks of HFD feeding. Right, AUC was used to quantify the ITT results. n = 5 mice in each group. Data are presented as the mean ± SEM.Two-way ANOVA followed by multiple comparison tests was performed for body weight curves (B, C). One-way ANOVA followed by multiple comparison tests was performed in (D-I) and AUC values of GTT and ITT (J, K).

To determine whether the anti-obesity effect of QUE was associated with altered energy intake or intestinal energy loss, we further measured cumulative food intake, fecal output, and fecal energy content. As shown in Supplementary Figure 1A-C, QUE treatment did not markedly affect cumulative food intake, daily stool weight, or fecal energy content compared with the HFD group, suggesting that the reduction in body weight gain induced by QUE was unlikely to be attributable to decreased caloric intake or increased energy excretion.

Regarding lipid metabolism, QUE administration resulted in a significant reduction in serum levels of free fatty acids (FFA, Figure 1E), triglycerides (TG, Figure 1F), and hepatic triglyceride content (Figure 1G). Analysis of body composition indicated that QUE significantly decreased fat mass while increasing lean mass (Figure 1H), suggesting a beneficial alteration in body composition. In terms of glucose homeostasis, QUE treatment significantly reduced fasting blood glucose levels (Figure 1I). In contrast, serum insulin levels were not obviously altered between the HFD and HFD+QUE groups (Supplementary Figure 1D). The glucose tolerance test (GTT) and insulin tolerance test (ITT) further revealed that QUE significantly enhanced glucose tolerance and insulin sensitivity. Notably, the blood glucose excursion during the GTT was markedly reduced in the QUE group, as evidenced by a significantly lower area under the curve (AUC, Figure 1J). Similarly, QUE-treated mice exhibited a more pronounced hypoglycemic response to insulin, accompanied by a significantly lower AUC in the ITT (Figure 1K).

Taken together, these findings indicate that QUE, particularly at a dose of 50 mg/kg, effectively suppresses HFD-induced body weight gain and fat accumulation, without significantly affecting food intake or fecal energy loss. Moreover, QUE alleviates HFD-associated metabolic disorders by improving lipid metabolism, reducing adiposity, and enhancing glucose tolerance and insulin sensitivity.

### Result 2: QUE boosts thermogenesis, increases energy expenditure, and reduces tissue damage in obesity caused by a high-fat diet

In the obesity model induced by a HFD, we conducted a comprehensive evaluation of the effects of QUE on thermogenic capacity, energy metabolism, and histopathology. Hematoxylin and eosin (H&E) staining revealed that, in comparison to the HFD group, the BAT in the HFD+QUE group exhibited a more pronounced multilocular lipid droplet phenotype. Additionally, there was a reduction in adipocyte size and lipid accumulation in both iWAT and eWAT, suggesting that QUE mitigates the morphological abnormalities in adipose tissue induced by a HFD (Figure 2A). Metabolic cage analysis demonstrated that QUE treatment significantly enhanced oxygen consumption (VO_2_), carbon dioxide production (VCO_2_), and heat production (HEAT), with these effects being evident throughout the light – dark cycle and more pronounced during the dark phase. The overall respiratory exchange ratio (RER) showed no significant difference, indicating an absence of a substantial shift in substrate utilization (Figures 2B-E). Furthermore, QUE significantly decreased serum ALT and AST levels, suggesting that QUE mitigates liver injury and hepatic dysfunction associated with a HFD (Figures 2F-G). Mechanistically, Western blot analysis revealed that QUE substantially upregulated UCP1 protein expression in BAT, while Ucp1 expression in iWAT exhibited an upward trend without reaching statistical significance (Figures 2H-I). In conclusion, QUE enhances energy expenditure primarily through the promotion of Ucp1-mediated thermogenesis in brown adipose tissue, thereby ameliorating metabolic disorders and mitigating histopathological damage induced by a HFD.

**Figure 2.**
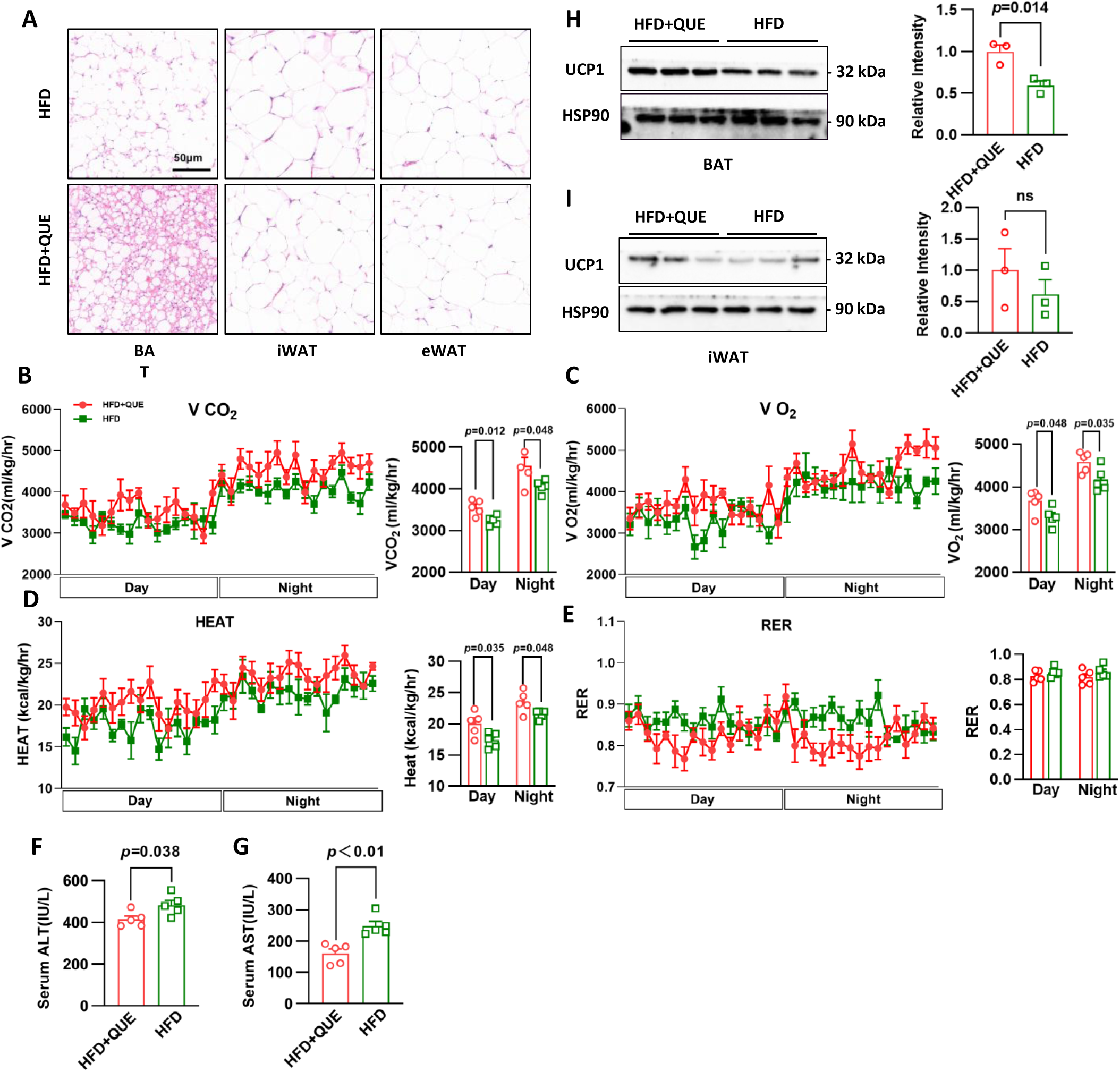
In high-fat diet-induced obesity, QUE enhances thermogenesis and energy expenditure by upregulating UCP1 expression, and alleviates histopathological damage. Hematoxylin and eosin staining of the BAT, iWAT and eWAT of mice in figure1. Scale bar: 50 μm. (B – E) Whole-body VO_2_, VCO_2_, heat generation,ambulatory activity, and RER were measured by metabolic cages of HFD and HFD+QUE mice after 9 weeks of HFD feeding. n = 5 mice in each group. Right panelpresents the statistical analyses of the data from the left panel per 12-h light/dark cycle in HFD and HFD+QUE mice. (F-G) ALT and AST level of mice in figure1. n = 5 mice in each group. Data are presented as mean ± SEM. Two-way ANOVA followed by multiple comparison tests was performed in (B-E). One-way ANOVA followed by multiple comparison tests was performed in (F-I).

### Result 3:QUE boosts energy use and cold tolerance during cold exposure by activating the UCP1-driven thermogenic process in fat tissues

To further substantiate the role of QUE in promoting thermogenic responses, we developed an acute cold exposure model using mice maintained on a standard chow diet and administered QUE via intraperitoneal injection prior to cold exposure (Figure 3A). In comparison to the control group, QUE administration significantly mitigated the reduction in core body temperature during cold exposure, resulting in an overall higher body temperature trajectory, which implies that QUE enhances cold tolerance (Figure 3B). Concurrently, QUE significantly upregulated the mRNA expression of Ucp1 in BAT, iWAT, and eWAT, indicating that QUE induces the expression of thermogenesis-related genes in adipose tissues (Figure 3C). With respect to metabolic phenotypes, QUE treatment altered the energy metabolic profile during cold exposure. Blood glucose levels showed a significant difference between groups (Figure 3D), whereas serum insulin levels were not markedly changed by QUE treatment (Supplementary Figure 2A), indicating that the glucose-lowering effect observed under cold exposure was unlikely to be due to increased circulating insulin. In contrast, oxygen consumption during cold exposure was significantly increased in the QUE-treated group (Figure 3E). Indirect calorimetry further demonstrated that QUE increased VO₂, VCO₂, and heat production, while the RER remained largely unchanged, suggesting that QUE primarily enhances energy expenditure and thermogenesis (Figure 3F). Histological analysis revealed that BAT in the QUE-treated group exhibited an increased presence of multilocular lipid droplets, while white adipose tissue demonstrated more pronounced browning/beiging characteristics (Figure 3G). Mechanistically, Western blot analysis corroborated that QUE treatment significantly elevated UCP1 protein expression levels in both BAT and iWAT (Figures 3H–I). In alignment with these findings, quantitative PCR analysis indicated a general upregulation of various genes associated with thermogenesis and mitochondrial function, including Ucp1, Pgc1α, Prdm16, Elovl3, Cidea, Cox7a1, and Cox8b, in BAT and iWAT (Figures 3J–K). In conclusion, QUE markedly enhances the UCP1-mediated thermogenic program in adipose tissues, particularly in BAT and iWAT, under conditions of cold exposure, thereby promoting energy expenditure and improving cold tolerance. This results in distinct pro-thermogenic and pro-browning effects.

**Figure 3.**
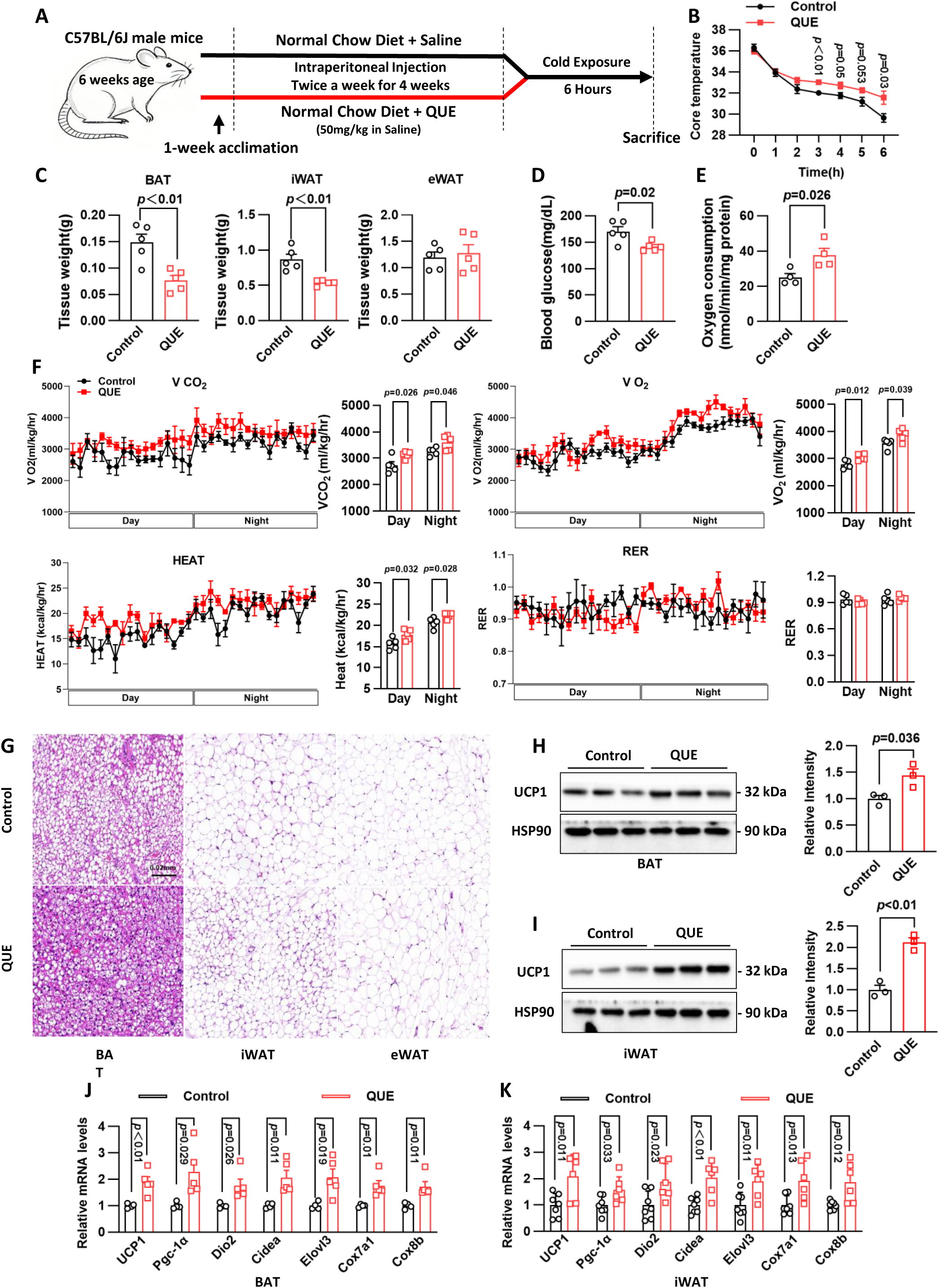
QUE enhances energy expenditure and cold tolerance under cold exposure by activating the UCP1-mediated thermogenic program in adipose tissues. C57BL/6J male mice fed on a Normal Chow Diet at 6 weeks of age and intraperitoneally injected with QUE(50 mg/kg) or saline twice a week for 4 weeks. Then mice were acutely exposed to 4 °C for 6 hours for other assays. Core temperature in mice of (A) during cold exposure. n =5 mice in each group. Tissue weight of BAT, iWAT, eWAT and Liver of mice of (A). n = 5 mice in each group. (D-E) Blood glucose and Oxygen consumption of mice of (A). n = 5 mice in each group. (F) Whole-body VO_2_, VCO_2_, heat generation,ambulatory activity, and RER were measured by metabolic cages of mice of (A). n = 5 mice in each group. Right panelpresents the statistical analyses of the data from the left panel per 12-h light/dark cycle. (G) Hematoxylin and eosin staining of the BAT, iWAT and eWAT of mice of (A). Scale bar: 0.02mm. (H-I) Left: UCP1 protein levels in the BAT(H) and iWAT(I) from mice in (A). Right: quantification of protein levels. n = 3 biological replicates. (J-K) The mRNA level of Ucp1, Pgc-1α, Dio2, Cidea, Elovl3, Cox7a1, and Cox8b in BAT(J) and iWAT(K) from mice in (A).n =5 mice in each group. Data are presented as mean ± SEM.Two-way ANOVA followed by multiple comparison tests was performed in (B) and (F). One-way ANOVA followed by multiple comparison tests was performed in (C-E) and (H-K).

### Result 4: QUE upregulates COX2 by stable binding and inhibiting proteasomal degradation, thereby activating the COX2–PGE2/EP signaling axis

Subsequently, a proteomic analysis was conducted on primary BAT cells derived from both the control and QUE-treated groups (Figure 4A). The analysis of differentially expressed proteins indicated significant alterations in multiple proteins post-QUE treatment. Among these, COX2 (Ptgs2) emerged as a pivotal differential protein and was consequently selected for further investigation (Figure 4B).

**Figure 4.**
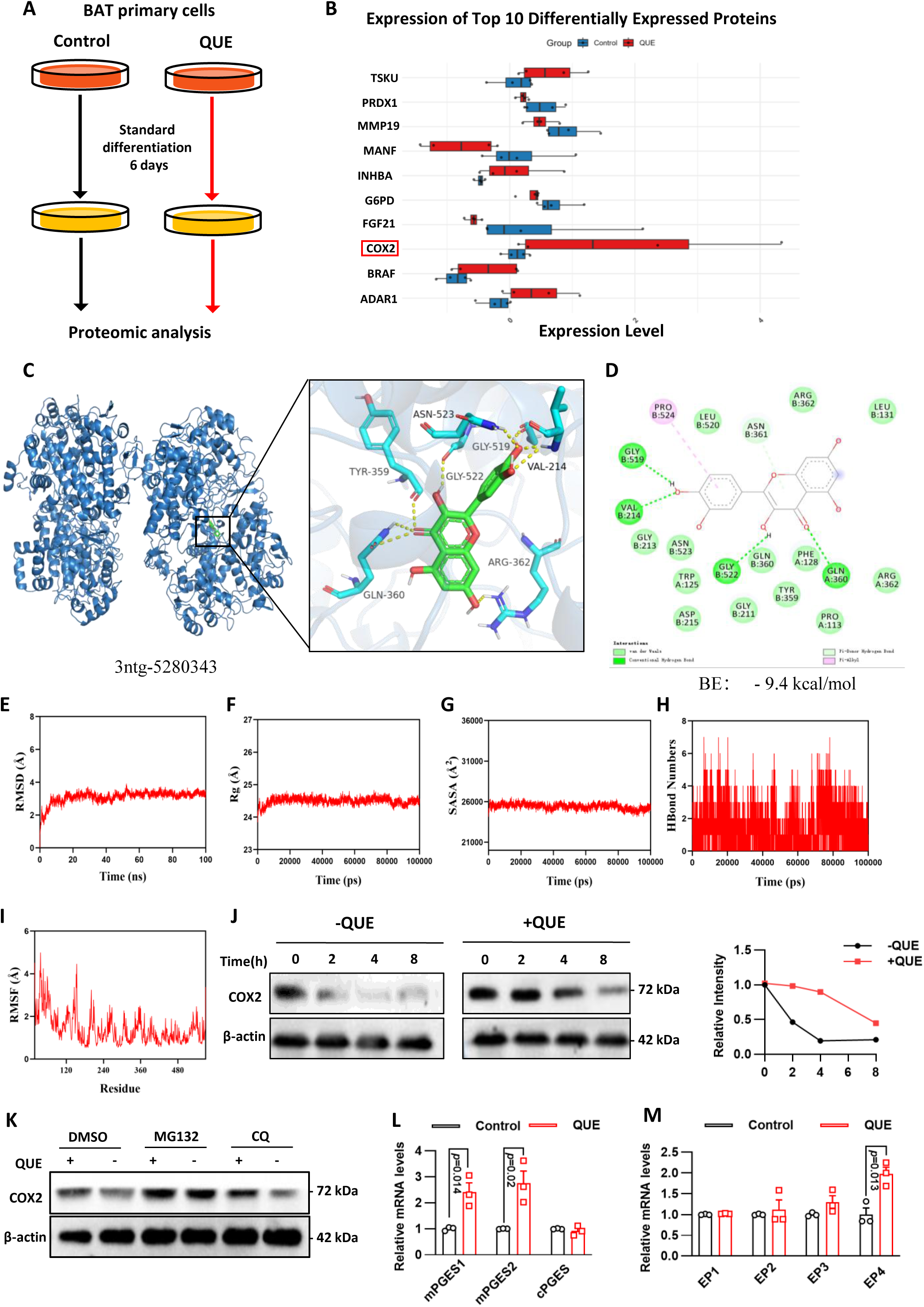
QUE upregulates COX2 by stable binding and inhibiting proteasomal degradation, thereby activating the COX2. – **PGE2/EP signaling axis** Primary BAT cells were isolated and subjected to standard differentiation for 6 days, followed by proteomic analysis in control and QUE-treated groups. Bar chart showing the expression of the top 10 differentially expressed proteins identified by proteomic analysis.Data are presented as mean ± SD. Three-dimensional binding model of QUE with COX2 (PDB: 3ntg-5280343). The detailed binding pocket and key interacting residues are highlighted in the magnified view. Two-dimensional interaction diagram showing the binding mode of QUE within the COX2 pocket. The predicted binding energy (BE) is −9.4 kcal/mol, with green circles indicating hydrogen bonds and other residues involved in van der Waals and hydrophobic interactions. Root-mean-square deviation (RMSD) of the QUE-COX2 complex over the entire MD simulation time course. Radius of gyration (Rg) analysis reflecting the compactness of the QUE-COX2 complex structure. Solvent-accessible surface area (SASA) of the QUE-COX2 complex during the simulation. Dynamic changes in the number of hydrogen bonds formed between QUE and COX2 residues over the simulation period. The RMSF profile of individual amino acid residues in the COX2 protein during MD simulation. Left: Western blot analysis of COX2 protein levels in cells treated with or without QUE over a time course of 0, 2, 4, and 8 hours. β-actin was used as a loading control. Right: Quantitative analysis of relative COX2 protein intensity, normalized to the 0-hour time point. Western blot analysis of COX2 protein levels in cells treated with DMSO (vehicle control), the proteasome inhibitor MG132, or the lysosome inhibitor chloroquine (CQ), in the presence or absence of QUE. β-actin was used as a loading control. Relative mRNA levels of PGE2 synthases (mPGES1, mPGES2, cPGES) in control and QUE-treated cells. Data are presented as mean ± SEM. Relative mRNA levels of PGE2 receptors (EP1, EP2, EP3, EP4) in control and QUE-treated cells.Data are presented as mean ± SEM. One-way ANOVA followed by multiple comparison tests was performed in (K-M).

To investigate the potential direct interaction between QUE and cyclooxygenase-2 (COX2), a series of computational analyses were performed, integrating molecular docking and molecular dynamics (MD) simulations. The two-dimensional structure of the ligand was sourced from PubChem and subsequently converted to a three-dimensional conformation in mol2 format. A high-resolution crystal structure of COX2 was selected from the RCSB Protein Data Bank (PDB) to serve as the receptor. Preprocessing of the receptor, which included the removal of water molecules and the stripping of ligands and ions, was executed using PyMOL. This was followed by hydrogenation, torsion definition, and the establishment of docking box parameters in AutoDock Vina (version 1.5.6) to determine the optimal binding conformation. The docking results revealed a strong binding affinity between QUE and COX2, with a predicted binding energy of −9.4 kcal/mol (Figures 4C – 4D). This finding aligns with the principle that a lower binding energy signifies a higher affinity and a more stable conformation. Subsequent interaction analysis, visualized using PyMOL and Discovery Studio 2019, revealed that QUE established a stable interaction network with several residues within the binding pocket. Specifically, it formed hydrogen bonds with GLY519, VAL214, GLY522, and GLN360; van der Waals interactions with GLY213, ASN523, TRP125, ASP215, GLY211, TYR359, PHE128, PRO113, ARG362, LEU131, and LEU520; and hydrophobic interactions with PRO524. This binding mode is indicative of favorable geometric complementarity and interaction strength (Figures 4C – 4D).

To assess the dynamic stability of the complex, the simulation trajectories were systematically analyzed using root mean square deviation (RMSD), radius of gyration (Rg), solvent-accessible surface area (SASA), the number of hydrogen bonds, and root mean square fluctuation (RMSF). The complex system achieved equilibrium after approximately 10 nanoseconds, with the RMSD exhibiting minor fluctuations around 3.3 Å, indicative of overall stability following ligand binding (Figure 4E). The Rg gradually stabilized after initial minor fluctuations, suggesting conformational adjustments without significant unfolding or instability during the simulation (Figure 4F). The SASA remained largely unchanged, indicating that QUE binding induced minimal structural perturbation to the protein’s surface exposure (Figure 4G). Analysis of hydrogen bonds revealed that QUE and COX2 maintained between 0 and 7 hydrogen bonds, predominantly around 4 bonds throughout the simulation, thereby supporting the persistence and reliability of hydrogen bonding interactions (Figure 4H). Furthermore, the Root Mean Square Fluctuation (RMSF) analysis revealed that the majority of residues exhibited fluctuations below 4 Å, indicating low overall flexibility and thereby corroborating the high stability of the complex (Figure 4I). Taken together, these computational findings indicate that QUE is capable of forming a stable binding interaction with COX2, maintaining a robust interaction network.

To elucidate the molecular mechanism by which QUE stabilizes the COX2 protein, cells were subjected to treatment with the proteasome inhibitor MG132, the lysosome inhibitor chloroquine (CQ), or a vehicle control (DMSO), both in the presence and absence of QUE (Figure 4K). Western blot analysis revealed that QUE treatment substantially elevated COX2 protein levels under DMSO control conditions. The inhibition of the proteasome by MG132 resulted in a significant increase in COX2 expression even in the absence of QUE, suggesting that COX2 undergoes constitutive degradation via the proteasomal pathway. Conversely, the inhibition of lysosomal function by CQ exerted minimal effects on COX2 protein levels, showing no significant difference compared to the DMSO group. This indicates a negligible role of the lysosomal pathway in COX2 degradation under the experimental conditions employed. These findings indicate that QUE primarily stabilizes the COX2 protein by inhibiting proteasome-mediated degradation.

Considering that COX2 is a crucial upstream rate-limiting enzyme in the biosynthesis of prostaglandin E2 (PGE2), and that PGE2 functions as a pivotal effector in mediating COX2-dependent inhibition of adipogenesis and limitation of adipose tissue expansion, we conducted an analysis of the mRNA expression levels of three key PGE2 synthases: microsomal prostaglandin E synthase 1 (mPGES1), microsomal prostaglandin E synthase 2 (mPGES2), and cytosolic prostaglandin E synthase (cPGES). This analysis aimed to determine whether QUE-induced stabilization of COX2 further activates the PGE2 synthetic pathway (Figure 4L). Quantitative PCR results demonstrated that QUE significantly upregulated the transcription of mPGES1 and mPGES2, while the expression of cPGES remained unchanged. These findings suggest that QUE preferentially enhances the mPGES1/2-mediated PGE2 synthesis pathway, predominantly affecting the microsomal route.

Given that PGE2 mediates its anti-adipogenic effects by activating downstream signaling pathways, such as the PKA pathway, via specific EP receptors, we concurrently assessed the mRNA expression levels of EP1 – EP4 (Figure 4M). The findings indicated that QUE significantly upregulated the expression of EP4, while exhibiting no notable impact on EP1, EP2, or EP3. This suggests that QUE may preferentially augment signaling through the PGE2-EP4 axis, thereby enhancing cellular responsiveness to PGE2 and promoting downstream anti-adipogenic processes at the receptor level. In conclusion, QUE primarily stabilizes COX2 protein by inhibiting proteasome-mediated degradation, significantly upregulates mPGES1 and mPGES2, and specifically enhances EP4 receptor transcription. This coordinated activation of the COX2-PGE2 synthesis and receptor signaling axis results in anti-adipogenic effects and improved metabolic phenotypes.

### Result 5. QUE enhances thermogenic gene expression and cAMP/PGE2 production in brown adipocytes through the COX2/EP4 pathway

To assess the impact of QUE on thermogenesis in brown adipocytes, we initially evaluated the expression of key genes associated with thermogenesis and lipid metabolism in immortalized brown adipocytes and 3T3-L1 cells. Western blot analysis demonstrated that treatment with QUE significantly elevated the protein levels of UCP1, peroxisome proliferator-activated receptor gamma coactivator 1-alpha (PGC-1α), cyclooxygenase 2 (COX2), and prostaglandin E2 receptor 4 (EP4) in both cell lines (Figure 5A, E). In alignment with the protein expression findings, qRT-PCR corroborated that QUE substantially increased the mRNA levels of Ucp1, Pgc-1α, Dio2, Cidea, Elovl3, Cox7a1 and Cox8b (Figure 5B, F), suggesting an enhancement in thermogenic and mitochondrial function.

**Figure 5.**
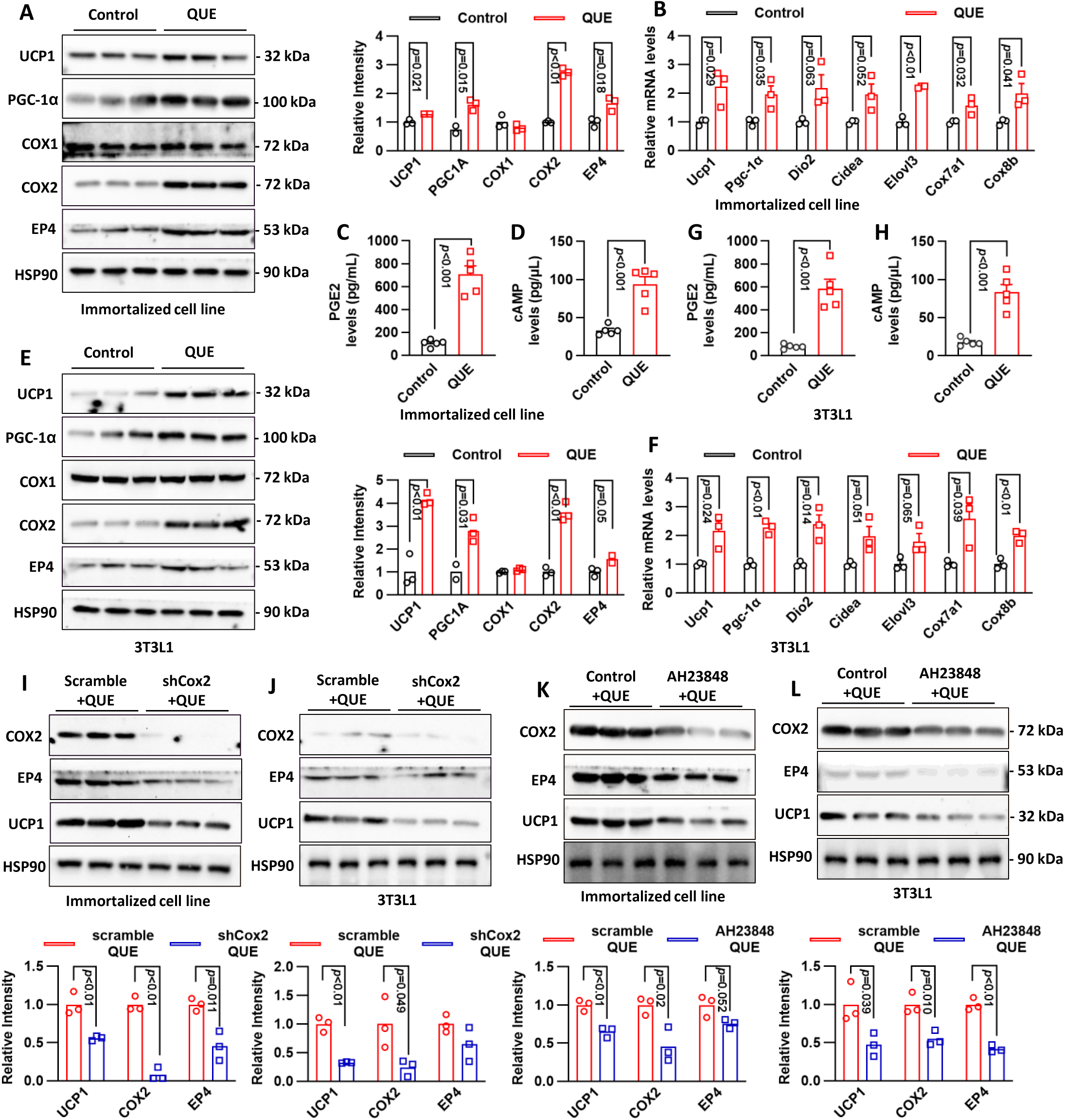
Quercetin (QUE) promotes thermogenic gene expression and cAMP/PGE2 production via the COX2/EP4 pathway in brown adipocytes. (A) Representative immunoblots and densitometric quantification of UCP1, PGC-1α, COX1, COX2, and EP4 protein levels in immortalized brown adipocytes treated with QUE or vehicle control. HSP90 was used as the loading control. (B) Relative mRNA levels of thermogenic genes, including Ucp1, Pgc1α, Dio2, Cidea, Elovl3, Cox7a1, and Cox8b, in immortalized brown adipocytes treated with QUE or vehicle control. (C, D) PGE2 (C) and cAMP (D) levels in immortalized brown adipocytes following QUE treatment. (E) Representative immunoblots and densitometric quantification of UCP1, PGC-1α, COX1, COX2, and EP4 protein levels in 3T3L1 cells treated with QUE or vehicle control. HSP90 was used as the loading control. (F) Relative mRNA levels of Ucp1, Pgc1α, Dio2, Cidea, Elovl3, Cox7a1, and Cox8b in 3T3L1 cells treated with QUE or vehicle control. (G, H) PGE2 (G) and cAMP (H) levels in 3T3L1 cells following QUE treatment. (I, J) Representative immunoblots and quantification of COX2, EP4, and UCP1 protein levels in immortalized brown adipocytes (I) and 3T3L1 cells (J) treated with QUE after Cox2 knockdown (shCox2) or scramble control. HSP90 was used as the loading control. (K, L) Representative immunoblots and quantification of COX2, EP4, and UCP1 protein levels in immortalized brown adipocytes (K) and 3T3L1 cells (L) treated with QUE in the presence or absence of the EP4 antagonist AH23848. HSP90 was used as the loading control. Data are presented as mean ± SEM. Each dot represents an individual sample. P values are indicated in the panels.

Additionally, treatment with QUE markedly increased the concentrations of prostaglandin E2 (PGE2) and cyclic adenosine monophosphate (cAMP) in both immortalized brown adipocytes (Figure 5C, D) and 3T3L1 cells (Figure 5G, H), indicating activation of the PGE2/cAMP signaling pathway. To verify the essential role of COX2 in facilitating the thermogenic effects of QUE, COX2 knockdown (shCox2) was conducted in both cell lines. Western blotting and densitometric analyses revealed that shCox2 negated the QUE-induced upregulation of COX2, EP4, and UCP1 protein expression (Figure 5I, J). Similarly, pharmacological inhibition of EP4 with AH23848 also impeded the QUE-mediated enhancement of COX2, EP4, and UCP1 levels (Figure 5K, L). These findings illustrate that QUE stimulates thermogenesis in brown adipocytes by activating the COX2/PGE2/EP4 signaling axis, which leads to increased cAMP production and the subsequent upregulation of thermogenic genes.

### Result 6: Adipocyte-specific Cox2 knockout (AdKO) eliminates QUE-induced thermogenesis in mice

To assess the in vivo significance of the COX2/EP4 pathway in QUE-mediated thermogenesis, we developed adipocyte-specific Cox2 AdKO mice and administered a 4-week regimen of QUE treatment (50 mg/kg, intraperitoneal injection, twice weekly), followed by 6 hours of cold exposure (refer to Figure 6A). Initially, we evaluated tissue weights and observed that QUE treatment significantly decreased the weights of iWAT, eWAT and BAT in control mice. However, this effect was completely absent in AdKO mice (see Figure 6B). Correspondingly, QUE treatment resulted in elevated blood glucose levels in control mice, whereas this increase was attenuated in AdKO mice (refer to Figure 6C). Furthermore, the QUE-induced enhancement in oxygen consumption, an indicator of thermogenic activity, was also nullified in AdKO mice (see Figure 6D).

**Figure 6.**
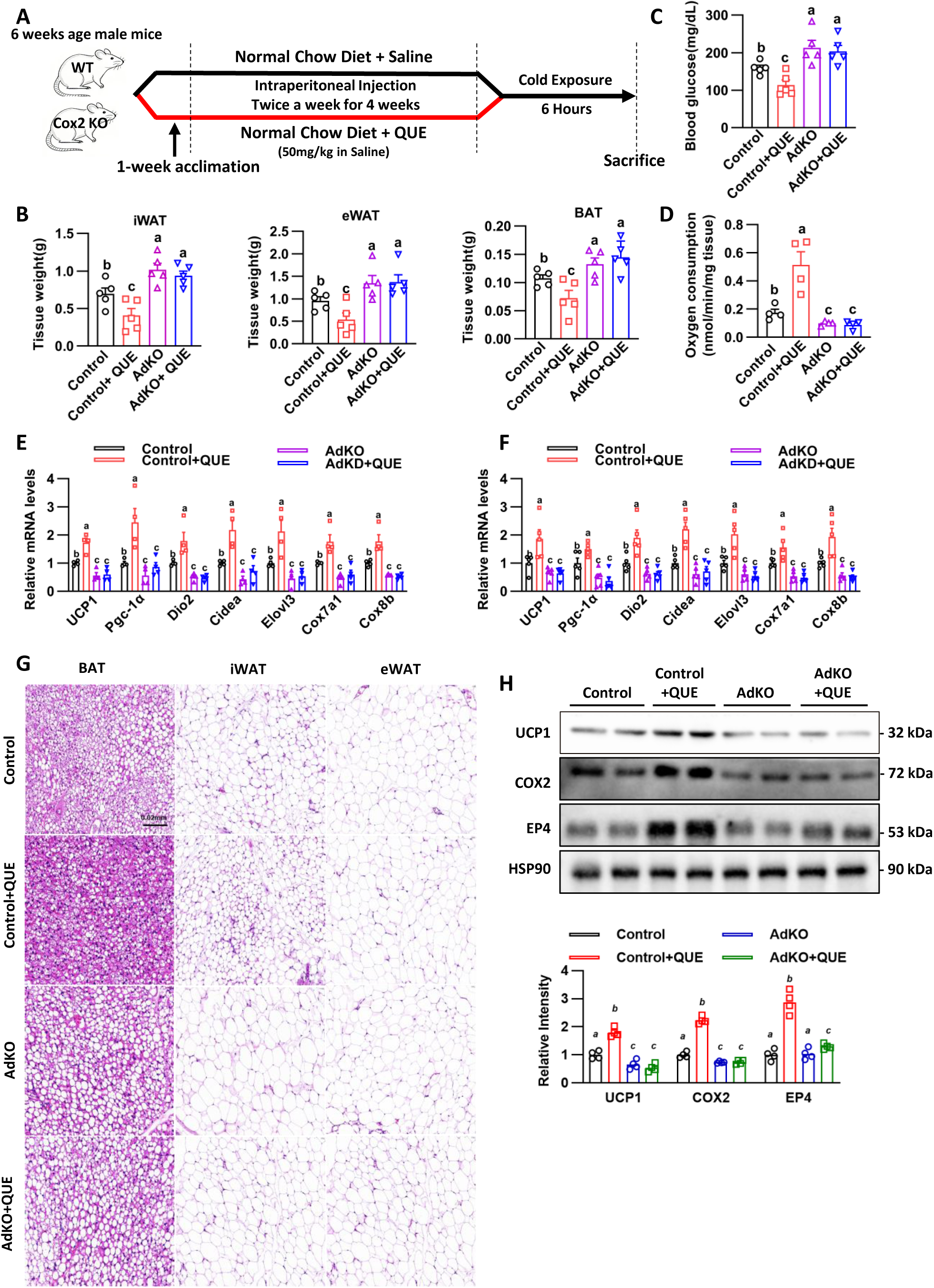
Adipocyte-specific Cox2 knockout (AdKO) abolishes quercetin (QUE)-induced thermogenesis in mice. (A) Schematic illustration of the experimental design. Six-week-old male control and adipocyte-specific Cox2 knockout (AdKO) mice were acclimated for 1 week, intraperitoneally injected with saline or QUE (50 mg/kg in saline) twice per week for 4 weeks under normal chow diet, followed by cold exposure for 6 h before sacrifice. (B) Weights of inguinal white adipose tissue (iWAT), epididymal white adipose tissue (eWAT), and brown adipose tissue (BAT) in the indicated groups. (C) Blood glucose levels in the indicated groups after cold exposure. (D) Oxygen consumption in the indicated groups. (E, F) Relative mRNA expression levels of thermogenic genes, including Ucp1, Pgc1α, Dio2, Cidea, Elovl3, Cox7a1, and Cox8b, in adipose tissues from the indicated groups. (G) Representative H&E staining images of BAT, iWAT, and eWAT from the indicated groups. (H) Representative immunoblots and densitometric quantification of UCP1, COX2, and EP4 protein levels in adipose tissue from the indicated groups. HSP90 was used as the loading control. Data are presented as mean ± SEM. Different lowercase letters indicate statistically significant differences among groups.

At the molecular level, qRT-PCR analysis demonstrated that QUE significantly upregulated the mRNA expression of thermogenic genes, including Ucp1, Pgc-1α, Dio2, Cidea, Elovl3, Cox7a1, and Cox8b, in BAT (Figure 6E) and iWAT (Figure 6F) of control mice. In contrast, these gene expression inductions were absent in adipocyte-specific Cox2 AdKO mice. Histological examination using hematoxylin and eosin (H&E) staining revealed that QUE treatment resulted in smaller lipid droplets in the BAT and iWAT of control mice, suggesting enhanced lipolysis and thermogenesis; however, this morphological change was not observed in AdKO mice (Figure 6G). Western blot analysis further corroborated that QUE elevated the protein levels of UCP1, COX2, and EP4 in control mice, whereas these effects were completely inhibited in AdKO mice (Figure 6H). Collectively, these in vivo findings indicate that the adipocyte-specific knockout of Cox2 negates the thermogenic and metabolic advantages conferred by QUE, thereby confirming the critical role of the COX2/EP4 signaling pathway in QUE-induced thermogenesis in brown adipocytes in mice.

## Discussion

In the present study, we systematically demonstrated that QUE)confers significant anti-obesity and metabolic benefits in models of high-fat diet (HFD)-induced obesity and acute cold exposure. Furthermore, we elucidated the COX2-PGE2/EP4 axis as a pivotal mechanism underlying its thermogenic effects. Our findings revealed that QUE significantly inhibited HFD-induced body weight gain, attenuated adipose tissue expansion and hepatic lipid accumulation, decreased both circulating and hepatic lipid levels, and enhanced glucose tolerance and insulin sensitivity. These results suggest that QUE effectively ameliorates obesity-associated glucose-lipid metabolic disorders. Notably, the beneficial metabolic effects of QUE were accompanied by increased energy expenditure and activation of adipose thermogenesis, indicating that its anti-obesity action is not solely attributable to reduced lipid storage but is also closely linked to augmented energy dissipation^14^.

A key finding of this study is the facilitation of thermogenesis by QUE in adipose tissues, particularly in BAT, and the induction of a browning-like phenotype in white adipose depots under appropriate physiological conditions^15^. In the HFD model, QUE significantly increased oxygen consumption, carbon dioxide production, and heat generation, while markedly upregulating the expression of UCP1 in BAT. Histological analyses revealed a more multilocular lipid droplet morphology in BAT and a reduction in adipocyte hypertrophy in iWAT and eWAT, suggesting an overall enhancement in adipose tissue function^16^. In the acute cold exposure model, QUE further improved cold tolerance, mitigated the decrease in core body temperature, and robustly induced the expression of thermogenic genes in BAT, iWAT, and eWAT.The results indicate that QUE not only reinstates thermogenic capacity in obese conditions but also augments adaptive thermogenic responses during cold exposure^17^. Notably, the thermogenic response was most pronounced in BAT, whereas the induction of browning-related characteristics in white adipose tissue was more evident under cold exposure^18^. This implies that the tissue-specific and context-dependent effects of QUE may be modulated by the physiological state.

Our study elucidates the mechanistic pathways through which QUE induces adipose thermogenesis, specifically by stabilizing COX2 and subsequently activating the COX2-PGE2-EP4 signaling cascade. Proteomic analyses have identified COX2 as a significantly differentially expressed protein in QUE-treated primary brown adipocytes. Further insights from molecular docking and molecular dynamics simulations indicate that QUE can stably bind to COX2, characterized by favorable binding energy and sustained intermolecular interactions, suggesting a potential direct interaction between QUE and COX2. Although computational evidence alone is insufficient to definitively confirm direct molecular binding within a biological context, these findings provide a robust structural framework for subsequent mechanistic investigations. Notably, our protein degradation assays revealed that QUE enhances COX2 protein abundance predominantly by inhibiting proteasome-mediated degradation, as opposed to lysosomal degradation. This observation is particularly significant as it implies that QUE modulates COX2 not only at the transcriptional level but also through post-translational stabilization, thereby extending the functional availability of this critical enzyme.

Considering that COX2 serves as the rate-limiting enzyme in the biosynthesis of PGE2^19^, we conducted an in-depth investigation to determine whether QUE-induced stabilization of COX2 leads to the activation of downstream prostaglandin signaling pathways. Our findings reveal that QUE significantly upregulates the expression of mPGES1 and mPGES2, while not affecting cPGES, thereby suggesting a preferential activation of the microsomal PGE2 synthesis pathway. Concurrently, QUE selectively enhances EP4 expression without significantly impacting EP1, EP2, or EP3, indicating that the downstream effects of QUE are likely mediated predominantly through the PGE2-EP4 axis. In alignment with these observations, QUE markedly elevates both PGE2 and cAMP levels in immortalized brown adipocytes and 3T3L1 cells, accompanied by a robust upregulation of thermogenic markers such as UCP1, PGC-1α, Dio2, Cidea, Elovl3, Cox7a1, and Cox8b. Given that cAMP functions as a central second messenger in canonical thermogenic signaling^20,21^, these results support a model wherein QUE stabilizes COX2, enhances PGE2 production, activates EP4-dependent signaling, increases intracellular cAMP, and ultimately induces a UCP1-driven thermogenic program^22,23^.

A notable strength of this study lies in the validation of the functional significance of the COX2/EP4 pathway through both pharmacological and genetic loss-of-function methodologies. In vitro analyses demonstrated that the knockdown of Cox2 completely eliminated the QUE-induced upregulation of COX2, EP4, and UCP1. Similarly, pharmacological inhibition of EP4 using AH23848 effectively obstructed the thermogenic effect of QUE. These findings suggest that the COX2/EP4 pathway is not merely associated with QUE’s mechanism of action but is essential for its thermogenic activity. Furthermore, in adipocyte-specific Cox2 knockout mice, the advantageous effects of QUE on adipose tissue mass, oxygen consumption, histological remodeling, and the expression of thermogenic genes and proteins were substantially diminished. These in vivo results underscore the role of adipocyte COX2 as a critical mediator of QUE-induced thermogenesis and affirm the physiological relevance of the COX2/EP4 signaling axis in this context.

From a translational standpoint, our findings indicate that QUE may serve as a promising natural compound for addressing obesity and related metabolic disorders by enhancing energy expenditure, rather than merely focusing on reducing caloric intake or lipid synthesis. This mechanism is particularly appealing, as strategies that enhance BAT activity and promote the browning of white adipose tissue have emerged as significant therapeutic approaches for obesity management^24–27^. Beyond its metabolic advantages, QUE also decreased serum ALT and AST levels, suggesting a protective effect against HFD-induced liver injury and hepatic dysfunction^28^. Consequently, QUE may provide broader systemic benefits by simultaneously improving adipose tissue function, hepatic lipid metabolism, and overall metabolic homeostasis.

This study has several limitations that should be acknowledged. Firstly, all in vivo experiments were conducted exclusively on male mice, leaving the potential influence of sex differences on the metabolic and thermogenic responses to QUE undetermined^29^. Secondly, QUE was primarily administered via intraperitoneal injection in this study, whereas oral administration would be more pertinent for nutritional or clinical applications. Consequently, further investigation is needed to assess the bioavailability, metabolism, and tissue distribution of QUE when delivered through physiologically relevant routes. Thirdly, although our data underscore the central role of adipocyte COX2/EP4 signaling, we cannot rule out the involvement of other pathways, such as sympathetic activation^30–32^, mitochondrial remodeling^33,34^, inflammatory modulation^35,36^, or gut microbiota-related mechanisms^37^, in contributing to the metabolic effects of QUE. Lastly, given that prostaglandin signaling can have context-dependent effects across different tissues^38–41^, the long-term safety and systemic implications of sustained activation of the COX2-PGE2/EP4 pathway necessitate further evaluation.

In summary, this study provides evidence that QUE significantly mitigates HFD-induced obesity and systemic metabolic dysfunction, while simultaneously enhancing adaptive thermogenesis and cold tolerance. Mechanistically, QUE stabilizes COX2 protein by inhibiting its proteasomal degradation, thereby activating the COX2-microsomal prostaglandin E synthase-1/2 (mPGES1/2)-prostaglandin E2 (PGE2)-EP4-cAMP signaling pathway and promoting a thermogenic program dependent on UCP1 in adipose tissues. The absence of QUE responsiveness in adipocyte-specific Cox2 knockout mice further corroborates the essential role of adipocyte COX2 in mediating its thermogenic and metabolic effects. Collectively, these findings not only enhance the current understanding of the metabolic actions of QUE but also provide a mechanistic foundation for the development of thermogenesis-targeted natural interventions against obesity and metabolic disorders.

## Conclusion

Our study establishes that quercetin acts as a potent activator of adipose thermogenesis, effectively ameliorating high-fat diet-induced obesity and enhancing cold tolerance. Mechanistically, we demonstrate that quercetin stabilizes COX2 protein by inhibiting its proteasomal degradation, thereby triggering the COX2-PGE2-EP4-cAMP signaling cascade and driving UCP1-dependent thermogenic programming. Consequently, targeting the COX2-mediated prostaglandin pathway with quercetin offers a promising nutritional strategy to combat metabolic disorders by shifting the energy balance towards increased expenditure rather than mere storage.

## Materials and Methods

### Animal studies

All animal procedures were performed in accordance with the Guidelines for Care and Use of Laboratory Animals of Binzhou Medical University Hospital and approved by the Animal Ethics Committee of Binzhou Medical University Hospital. Male C57BL6 mice aged 6-8 weeks were purchased from Jinan Pengyue Laboratory Animal Breeding Co., Ltd, Jinan, China. Mice were housed in a 12-hour light and 12-hour dark cycle at 22 °C with free access to water and a standard rodent chow diet unless otherwise specified.

A high-fat diet (HFD) with 60% of calories from fat was purchased from Research Diets (D12492). After one week of adaptive feeding, six-week-old mice were administered a high-fat diet and concurrently received intraperitoneal injections of quercitrin (10, 50, and 100 mg/kg) twice a week for 9 weeks, during which body weight and other parameters were monitored.

A glucose tolerance test (GTT) and an insulin tolerance test (ITT) were performed in mice after a HFD. Mice fasted for 16 hours (GTT) or 5 hours (ITT) following intraperitoneal injection with glucose (2 g per kg body mass) or insulin (0.75 U per kg body mass), respectively. For indirect calorimetry assay, mice were placed in the Columbus Instruments Comprehensive Lab Animal Monitoring System and allowed to acclimatize. The energy expenditure and respiration were monitored at the basal state.

For acute cold exposure, male C57BL/6J mice housed at 22 °C were transferred to 4 °C for the time indicated, and their rectal temperature was monitored every hour using the animal electronic thermometer (ALT-ET03, Shanghai Alcott Biotech Co. Ltd).

### Histological analysis

Dissected adipose tissues were fixed in 10% neutral buffered formalin. Hematoxylin and eosin (H&E) staining was performed by Pinuofei Biotechnology (Wuhan, China).

### Serum insulin, triglyceride, PEG2 and cAMP levels

The serum insulin level was measured through an insulin ELISA kit. The triglyceride level was analyzed using a triglyceride assay kit according to the manu facturer’s instructions. The cellular PGE2 level was measured using a PGE2 ELISA kit. Similarly, the intracellular cAMP level was determined by a cAMP ELISA kit following the manufacturer’s protocol.

### Cell culture and differentiation

Immortalized brown preadipocytes were generated, cultured and differentiated as previously described ^42^. Briefly, preadipocytes were cultured in DMEM supplemented with 10% fetal bovine serum (FBS), 100 U/ml penicillin and 0.1 mg/ml streptomycin (Strep). Maintenance medium (MM) was prepared by adding 20 nM insulin and 1 nM triiodo-L-thyronine to the culture medium. Induction medium (IM) was prepared by adding 0.5 µM dexamethasone, 0.5 mM isobutylmethylxanthine (IBMX) and 0.125 mM indomethacin to MM. When the cell density reached 70% (day -2), the cells were cultured in MM. Upon reaching 100% confluence (day 0), the medium was switched to IM and the cells were induced for 48 h (day 0-2). Then the cells were maintained in MM for an additional 4 days (day 2-6) to allow for complete differentiation.

Preadipocytes 3T3L1 were cultured in DMEM containing 10% Newborn Calf Serum (NBCS), 1% Biotin, pantothenate and 1% penicillin-streptomycin. Two days after reaching 100% confluence (designated day 0), preadipocyte 3T3L1 were induced to differentiate into white adipocytes with induction medium (DMEM containing 10% FBS, 0.5 mM 3-isobutyl-1-methylxanthine, 1 µM Dexamethasone and 170 nM insulin) until day 2. Cells were then cultured in DMEM containing 10% FBS and 170 nM Insulin for 2 days. From days 4 to 8, cells were cultured with DMEM containing 10% FBS, which was changed every other day. Usually, the cells were harvested on day 8 for subsequent assays.

All cells were kept in a 5% CO2 incubator at 37℃, and were tested negative for mycoplasma before the experiments.

### RNA preparation and quantitative real-time PCR

Total RNA was extracted from the cells and tissues using RNA Extraction Reagent, and reverse transcription was performed with HiScript III RT SuperMix for qPCR (+gDNA wiper). Quantitative real-time PCR was performed using ChamQ Universal SYBR Qpcr Master Mix in a thermal cycler (Applied Biosystems, CA, USA). Relative mRNA expression was determined by the ΔΔCt method. The sequences of the qPCR primers used in this study are listed in Table S1.

### Protein extraction and Western blotting

Total protein was extracted from cells or adipose tissues using lysis buffer [100 mM NaCl, 50 mM Tris (pH 7.5), 0.5% Triton X-100, 5% glycerol] supplemented with phosphatase inhibitor cocktail and phenylmethylsulfonyl fluoride (PMSF). Lysates were run on 10% SDS-PAGE gels and immunoblotted with primary antibodies against HSP90, UCP1, COX2, EP4, PGC1α, and β-actin. HRP-conjugated secondary antibodies were used (anti-rabbit, anti-mouse). The intensity of bands was quantified using ImageJ, and the values of target proteins were normalized to those of the internal control on the same membrane. The antibodies used in this study are summarized in Table S2, while the other reagents are listed in Table S3.

### Isolation of Mature Adipocytes and Oxygen Consumption Measurement

Approximately 50 mg of fresh animal adipose tissue was rapidly excised and minced into fine pieces (1-2 mm³) and transferred to a digestion solution containing 1 mg/ml Collagenase Type II in DMEM. The mixture was incubated at 37°C for 40 min. During incubation, the tissue was gently triturated every 10 min to facilitate dispersion.

The digestion was terminated by adding DMEM supplemented with 10% Fetal Bovine Serum (FBS). The suspension was filtered through a 100 μm mesh to remove undigested fragments. To isolate mature adipocytes, the filtrate was centrifuged at 200×g for 5 min. Due to their buoyancy, the upper layer (containing isolated mature adipocytes) was carefully collected.

For the assay, the isolated adipocytes were washed gently and resuspended in 1 ml oxygen-saturated buffer (phosphate-buffered saline supplemented with 25 mM glucose, 2% BSA and 1 mM pyruvate). Oxygen consumption was measured using a Clark electrode (Oxygraph+ system, Hanstech, UK). Data were normalized to total protein content.

### Plasmids and viruses

The short hairpin RNA (shRNA) lentiviral plasmid was cloned into pSP-108 vector (Addgene, MA, USA) and co-transfected into HEK293T cells together with VSV-G plasmid (Addgene, MA, USA) and psPAX2 plasmid (Addgene, MA, USA) for virus production. The viral supernatant was collected 48 hours after transfection and concentrated using PEG 8000.

Immortalized brown preadipocytes and 3T3L1 cells were infected with the knockdown lentivirus, selected with puromycin, and plated for differentiation.

### Statistical analysis

Statistical analyses were performed using GraphPad Prism 10.0 software. Data were presented as mean ± standard error of the mean (SEM). Statistical differences between two groups were assessed using unpaired two-tailed Student’ s t-test, and one-way or two-way ANOVA as indicated. Statistical significance was defined as P < 0.05.

## Data availability

All data, methods, and results of statistical analyses are reported in this paper and the associated Supplementary Materials. Any specific inquiries can be addressed to the corresponding author.

## Ethics and consent to participate

All animal experiments were approved and complied with the guidelines of the Institutional Animal Care and Use Committee of the Binzhou Medical University Hospital, China (permit number: 20211008-24).

## Consent for publication

Not applicable.

## Availability of data and materials

Not applicable.

## Acknowledgements

This work was supported by the Special Funds of Taishan Scholars Project of Shandong Province (grant number: tsqn 202312384).

---

**Supplementary Figure 1.**
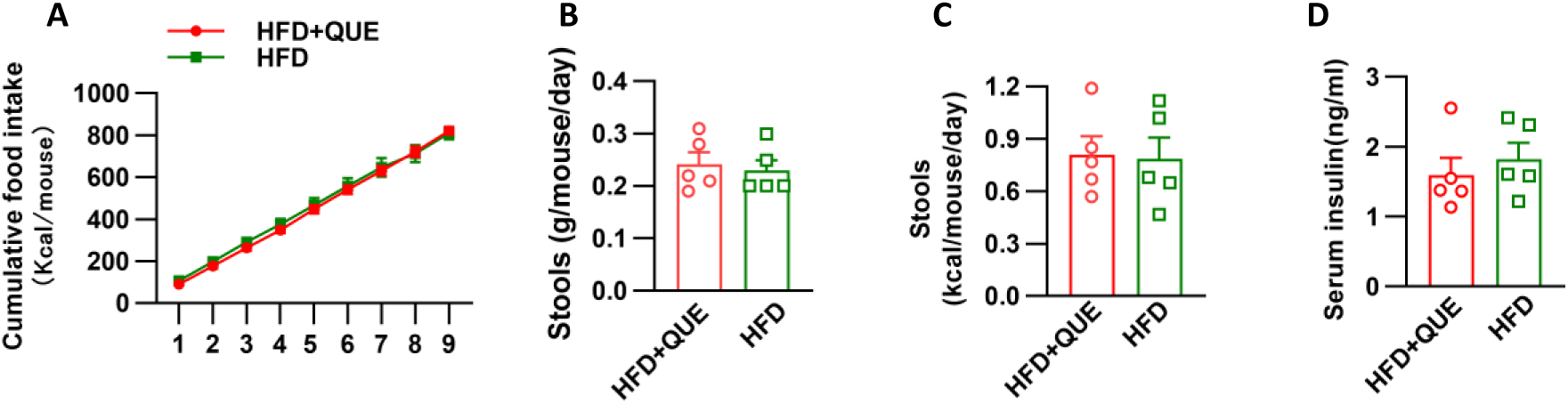
Effects of QUE on food intake, fecal output, and serum insulin levels in HFD-fed mice. (A) Cumulative food intake of mice fed a high-fat diet (HFD) and treated with QUE or vehicle during the experimental period. (B) Fecal output (g/mouse/day) in HFD-fed mice with or without QUE treatment. (C) Fecal energy content (kcal stools/day) in HFD-fed mice with or without QUE treatment. (D) Serum insulin levels in HFD-fed mice with or without QUE treatment. Data are presented as mean ± SEM. Each symbol represents one mouse.

**Supplementary Figure 2.**
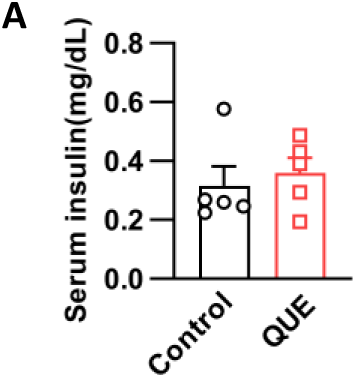
Serum insulin levels in control and QUE-treated mice. (A) Serum insulin levels in the Control and QUE groups. Data are presented as mean ± SEM. Each symbol represents one mouse.

**Table S1.**
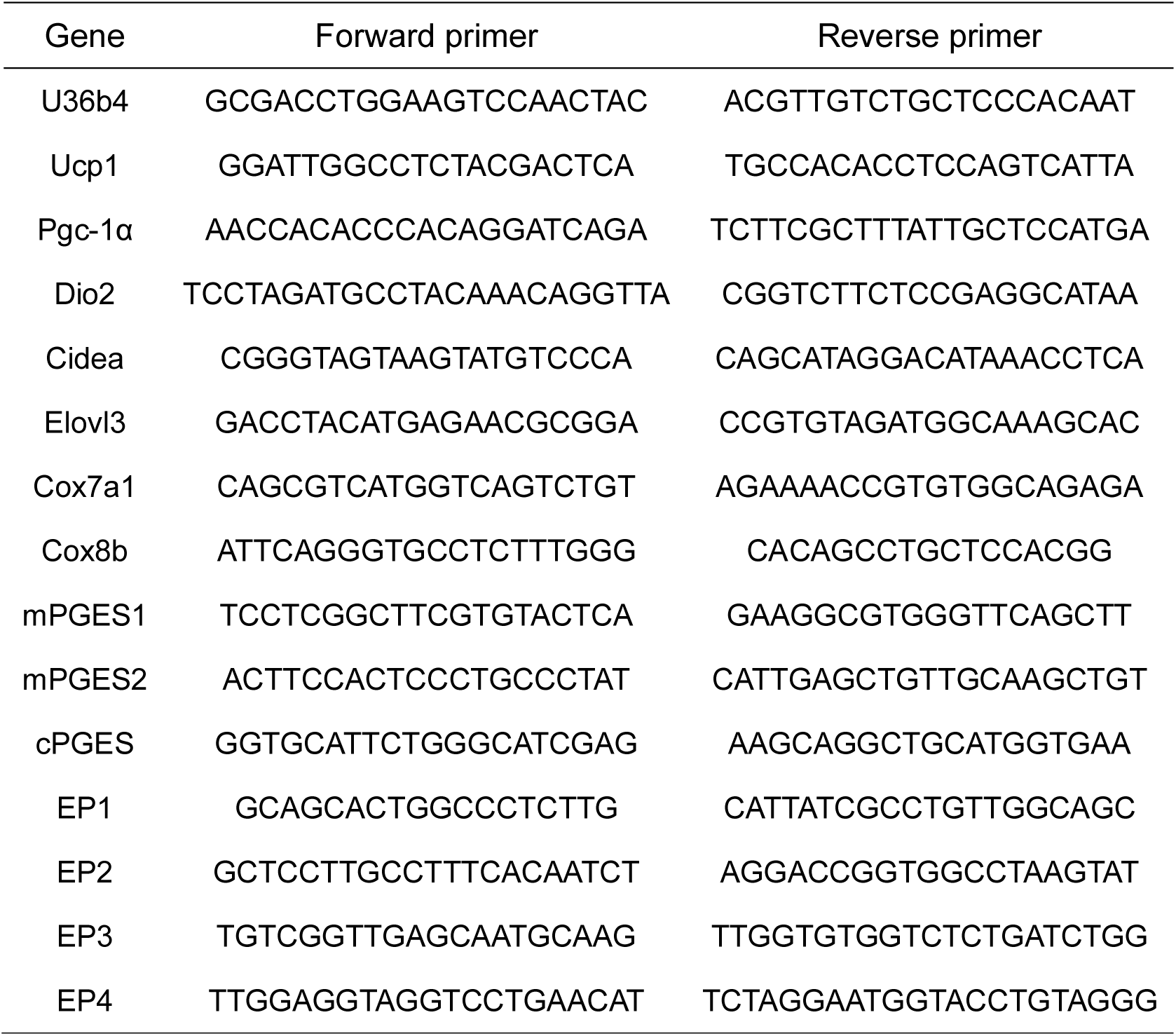
Quantitative PCR primers.

**Table S2.** Antibody.

| Company | Antibody | Product Code | Dilution ratio |
| --- | --- | --- | --- |
| Proteintech | HSP90 Polyclonal antibody | 13171-1-AP | 1:3000 |
| Alpha Diagnostic | UCP1 antibody | UCP11-A | 1:1000 |
| CST | Cox2 Antibody | 4842 | 1:1000 |
| Proteintech | PTGER4 Polyclonal antibody | 24895-1-AP | 1:1000 |
| Proteintech | PGC1a Monoclonal antibody | 66369-1-Ig | 1:1000 |
| Proteintech | Beta Actin Polyclonal antibody | 20536-1-AP | 1:1000 |
| Proteintech | HRP-conjugated Goat<br>Anti-Mouse IgG(H+L) | SA00001-1 | 1:10000 |
| Proteintech | HRP-conjugated Goat<br>Anti-Rabbit IgG(H+L) | SA00001-2 | 1:10000 |

**Table S3.** Other reagents.

| Company | Product Name | Product Code |
| --- | --- | --- |
| Rearch Diets | Rodent Diet With 60 kcal% Fat | D12492 |
| Pufei De<br>Biotech | Quercitrin | CAS: 522-12-3 |
| Mercodia | Insulin ELISA | 10-1113-01 |
| APPLYGEN | triglyceride assay kit | E1003-125 |
| Elabscience | PGE2 ELISA kit | E-EL-0034 |
| Sigma-Aldrich | cAMP ELISA kit | CA200-1KT |
| Meilunbio | DMEM | MA0212 |
| Hyclone | FETAL BOVINE SERUM CHARACTERIZED | SH30396.03 |
| Vazyme | ChamQ Universal SYBR Qpcr Master Mix | Q711 |
| Vazyme | RNA isolater Total RNA Extraction Reagent | R401 |
| Vazyme | HiScript III RT SuperMix for qPCR (+gDNA wiper) | R323-01 |
| Sigma-Aldrich | Insulin | I3536 |
| Sigma-Aldrich | T3 | T2877 |
| Sigma-Aldrich | IBMX | I5859 |
| Sigma-Aldrich | Indomethacin | I7378 |
| Sigma-Aldrich | Dexamethasone | D4902 |
| Meilunbio | Penicillin/Streptomycin,sterile (100X) | MA0110 |
| Hyclone | NBCS | SH30118.03 |
| Sigma-Aldrich | Biotin | #B4639 |
| Sinopharm<br>Chemical<br>Reagent Co.,<br>Ltd | pantothenate | #67000434 |
| MCE | AH23848 | HY-10419A |
| MCE | CQ | HY-17589A |
| MCE | MG-132 | HY-13259 |
| Yeasen | Inhibitor cocktail | 20141ES03 |
| Yeasen | PMSF | 58076ES |
| Sigma-Aldrich | Collagenase Type II | C2-BIOC |

## References

1. Dicker D, Karpati T, Promislow S, Reges O. Implications of the European Association for the Study of Obesity’s New Framework Definition of Obesity: Prevalence and Association With All-Cause Mortality. Ann Intern Med. Aug 2025;178(8):1065–1072. doi:10.7326/annals-24-02547

2. Conway B, Rene A. Obesity as a disease: no lightweight matter. Obes Rev. Aug 2004;5(3):145–51. doi:10.1111/j.1467-789X.2004.00144.x

3. Li L, Li B, Li M, Speakman JR. Switching on the furnace: Regulation of heat production in brown adipose tissue. Mol Aspects Med. Aug 2019;68:60–73. doi:10.1016/j.mam.2019.07.005

4. Sakers A, De Siqueira MK, Seale P, Villanueva CJ. Adipose-tissue plasticity in health and disease. Cell. Feb 3 2022;185(3):419–446. doi:10.1016/j.cell.2021.12.016

5. Alizadeh SR, Ebrahimzadeh MA. Quercetin derivatives: Drug design, development, and biological activities, a review. Eur J Med Chem. Feb 5 2022;229:114068. doi:10.1016/j.ejmech.2021.114068

6. Zhang L, Wang X, Chang L, et al. Quercetin improves diabetic kidney disease by inhibiting ferroptosis and regulating the Nrf2 in streptozotocin-induced diabetic rats. Ren Fail. Dec 2024;46(1):2327495. doi:10.1080/0886022x.2024.2327495

7. Jiang L, Yi R, Chen H, Wu S. Quercetin alleviates metabolic-associated fatty liver disease by tuning hepatic lipid metabolism, oxidative stress and inflammation. Anim Biotechnol. Dec 2025;36(1):2442351. doi:10.1080/10495398.2024.2442351

8. Yi R, Liu Y, Zhang X, et al. Unraveling Quercetin’s Potential: A Comprehensive Review of Its Properties and Mechanisms of Action, in Diabetes and Obesity Complications. Phytother Res. Dec 2024;38(12):5641–5656. doi:10.1002/ptr.8332

9. Okselni T, Septama AW, Juliadmi D, et al. Quercetin as a therapeutic agent for skin problems: a systematic review and meta-analysis on antioxidant effects, oxidative stress, inflammation, wound healing, hyperpigmentation, aging, and skin cancer. Naunyn Schmiedebergs Arch Pharmacol. May 2025;398(5):5011–5055. doi:10.1007/s00210-024-03722-3

10. Chiang MC, Tsai TY, Wang CJ. The Potential Benefits of Quercetin for Brain Health: A Review of Anti-Inflammatory and Neuroprotective Mechanisms. Int J Mol Sci. Mar 28 2023;24(7)doi:10.3390/ijms24076328

11. Zhang F, Zhang Y, Zhou J, et al. Metabolic effects of quercetin on inflammatory and autoimmune responses in rheumatoid arthritis are mediated through the inhibition of JAK1/STAT3/HIF-1α signaling. Mol Med. Oct 10 2024;30(1):170. doi:10.1186/s10020-024-00929-1

12. Qi W, Qi W, Xiong D, Long M. Quercetin: Its Antioxidant Mechanism, Antibacterial Properties and Potential Application in Prevention and Control of Toxipathy. Molecules. Oct 3 2022;27(19)doi:10.3390/molecules27196545

13. Cheng M, Yuan C, Ju Y, et al. Quercetin Attenuates Oxidative Stress and Apoptosis in Brain Tissue of APP/PS1 Double Transgenic AD Mice by Regulating Keap1/Nrf2/HO-1 Pathway to Improve Cognitive Impairment. Behav Neurol. 2024;2024:5698119. doi:10.1155/2024/5698119

14. Chondronikola M. The role of brown adipose tissue and the thermogenic adipocytes in glucose metabolism: recent advances and open questions. Current Opinion in Clinical Nutrition & Metabolic Care. 2020;23(4):282–287. doi:10.1097/mco.0000000000000662

15. Auger C, Kajimura S. Detouring adrenergic stimulation to induce adipose thermogenesis. Nature Reviews Endocrinology. 2021/10/01 2021;17(10):579–580. doi:10.1038/s41574-021-00546-6

16. Boudina S, Graham TE. Mitochondrial function/dysfunction in white adipose tissue. Exp Physiol. Sep 2014;99(9):1168–78. doi:10.1113/expphysiol.2014.081414

17. Auger C, Kajimura S. Detouring adrenergic stimulation to induce adipose thermogenesis. Nat Rev Endocrinol. Oct 2021;17(10):579–580. doi:10.1038/s41574-021-00546-6

18. Luo X, Jia R, Luo XQ, et al. Cold Exposure Differentially Stimulates Angiogenesis in BAT and WAT of Mice: Implication in Adrenergic Activation. Cell Physiol Biochem. 2017;42(3):974–986. doi:10.1159/000478680

19. Giuliano F, Warner TD. Origins of prostaglandin E2: involvements of cyclooxygenase (COX)-1 and COX-2 in human and rat systems. J Pharmacol Exp Ther. Dec 2002;303(3):1001–6. doi:10.1124/jpet.102.041244

20. Tabuchi C, Sul HS. Signaling Pathways Regulating Thermogenesis. Front Endocrinol (Lausanne). 2021;12:595020. doi:10.3389/fendo.2021.595020

21. Wen X, Zhang B, Wu B, et al. Signaling pathways in obesity: mechanisms and therapeutic interventions. Signal Transduct Target Ther. Aug 28 2022;7(1):298. doi:10.1038/s41392-022-01149-x

22. Xue K, Wu D, Wang Y, et al. The mitochondrial calcium uniporter engages UCP1 to form a thermoporter that promotes thermogenesis. Cell Metab. Sep 6 2022;34(9):1325–1341.e6. doi:10.1016/j.cmet.2022.07.011

23. St-Jacques B, Ma W. Peripheral prostaglandin E2 prolongs the sensitization of nociceptive dorsal root ganglion neurons possibly by facilitating the synthesis and anterograde axonal trafficking of EP4 receptors. Exp Neurol. Nov 2014;261:354–66. doi:10.1016/j.expneurol.2014.05.028

24. Prapaharan B, Lea M, Beaudry JL. Weighing in on the role of brown adipose tissue for treatment of obesity. J Pharm Pharm Sci. 2024;27:13157. doi:10.3389/jpps.2024.13157

25. Pan R, Zhu X, Maretich P, Chen Y. Combating Obesity With Thermogenic Fat: Current Challenges and Advancements. Front Endocrinol (Lausanne). 2020;11:185. doi:10.3389/fendo.2020.00185

26. Wang Q, Li D, Cao G, et al. IL-27 signalling promotes adipocyte thermogenesis and energy expenditure. Nature. Dec 2021;600(7888):314–318. doi:10.1038/s41586-021-04127-5

27. Lee YH, Jung YS, Choi D. Recent advance in brown adipose physiology and its therapeutic potential. Exp Mol Med. Feb 21 2014;46(2):e78. doi:10.1038/emm.2013.163

28. Im AR, Yang WK, Park YC, Kim SH, Chae S. Hepatoprotective Effects of Insect Extracts in an Animal Model of Nonalcoholic Fatty Liver Disease. Nutrients. Jun 7 2018;10(6)doi:10.3390/nu10060735

29. McDonald RB, Day C, Carlson K, Stern JS, Horwitz BA. Effect of age and gender on thermoregulation. Am J Physiol. Oct 1989;257(4 Pt 2):R700–4. doi:10.1152/ajpregu.1989.257.4.R700

30. Hu B, Lv X, Chen H, et al. Sensory nerves regulate mesenchymal stromal cell lineage commitment by tuning sympathetic tones. J Clin Invest. Jul 1 2020;130(7):3483–3498. doi:10.1172/jci131554

31. Guan J, Chen C, Wu S, Zhu H. The Role of PGE2 in Age-related Diseases. Curr Drug Targets. 2025;26(11):757–769. doi:10.2174/0113894501383329250616070727

32. Stone AJ, Copp SW, Kaufman MP. Role of prostaglandins in spinal transmission of the exercise pressor reflex in decerebrated rats. Neuroscience. Sep 26 2014;277:26–35. doi:10.1016/j.neuroscience.2014.06.061

33. Marques E, Burr SP, Casey AM, et al. An inherited mitochondrial DNA mutation remodels inflammatory cytokine responses in macrophages and in vivo in mice. Nat Commun. Nov 20 2025;16(1):10222. doi:10.1038/s41467-025-65023-4

34. Che L, Wu JS, Du ZB, et al. Targeting Mitochondrial COX-2 Enhances Chemosensitivity via Drp1-Dependent Remodeling of Mitochondrial Dynamics in Hepatocellular Carcinoma. Cancers (Basel). Feb 6 2022;14(3)doi:10.3390/cancers14030821

35. Adamczyk AM, Leicaj ML, Fabiano MP, et al. Extracellular vesicles from human plasma dampen inflammation and promote tissue repair functions in macrophages. J Extracell Vesicles. Jun 2023;12(6):e12331. doi:10.1002/jev2.12331

36. Wang Q, Xie Y, He Q, Geng Y, Xu J. LncRNA-Cox2 regulates macrophage polarization and inflammatory response through the CREB-C/EBPβ signaling pathway in septic mice. Int Immunopharmacol. Dec 2021;101(Pt B):108347. doi:10.1016/j.intimp.2021.108347

37. Zhang M, Wu X, Gao H, et al. Chinese Herbal Medicine for Irritable Bowel Syndrome: A Perspective of Local Immune Actions. Am J Chin Med. 2024;52(7):2079–2106. doi:10.1142/s0192415x24500800

38. Rahman MS. Prostacyclin: A major prostaglandin in the regulation of adipose tissue development. J Cell Physiol. Apr 2019;234(4):3254–3262. doi:10.1002/jcp.26932

39. Nakagawa T. Roles of prostaglandin E2 in the cochlea. Hear Res. Jun 2011;276(1-2):27–33. doi:10.1016/j.heares.2011.01.015

40. St Denis A, Simonette R, Rady PL, Tyring SK. The Role of Prostaglandin Pathway and EP Receptors in Skin Cancer Development. Int J Dermatol. Jul 2025;64(7):1186–1200. doi:10.1111/ijd.17711

41. Singh AK, Zajdel J, Mirrasekhian E, et al. Prostaglandin-mediated inhibition of serotonin signaling controls the affective component of inflammatory pain. J Clin Invest. Apr 3 2017;127(4):1370–1374. doi:10.1172/jci90678

42. Pan D, Fujimoto M, Lopes A, Wang YX. Twist-1 is a PPARdelta-inducible, negative-feedback regulator of PGC-1alpha in brown fat metabolism. Cell. Apr 3 2009;137(1):73–86. doi:10.1016/j.cell.2009.01.051

